# Rapid reductions in population size drive evolutionary divergence in diatoms

**DOI:** 10.1101/2022.09.25.509407

**Authors:** Nathan G. Walworth, Josh L. Espinoza, Phoebe A. Argyle, Jana Hinners, Naomi M. Levine, Martina A. Doblin, Chris L. Dupont, Sinéad Collins

## Abstract

Unicellular photosynthetic marine microbes, or phytoplankton, make up the base of marine food webs and drive global nutrient cycles. Despite their key roles in ecology and biogeochemistry, we have a limited understanding of how the basic features of their demographics along with dynamic environments affect trait evolution. A key feature of diatom ecology is frequent extreme reductions in population size, both as part of their bloom-and-bust growth dynamics, and as a result of living within ocean currents. Here, we use experimental evolution to understand which metabolic pathways and functions readily diversify in diatom populations following population bottleneck events. We subjected replicate populations of six genetically distinct diatom strains to population bottlenecks and then subsequently allowed them to evolve as large populations in the absence of environmental change. Phylogenies and global expression of orthologs were generally strain-specific, indicating that vertical (inherited) evolutionary constraints largely determine the occupation of specific locations in the transcriptional landscape (i.e. tran-scape). Following bottlenecks and subsequent evolution as large populations, transcriptional networks of most populations returned to those of the ancestral population. However, at least one replicate population per lineage migrated in the tran-scape, demonstrating that evolutionary changes in gene expression patterns and transcriptional relationships can be driven by population bottlenecks even in the absence of environmental change. Importantly, the orthologs dominating transcriptional diversification resided in common, central metabolic pathways. These data advance our understanding of constraints and patterns of transcriptional relationships underlying trait evolution in microbes that drive global food webs and elemental cycles.

## Introduction

Diatoms are among the most diverse and ubiquitous eukaryotic microbes in aquatic environments, yet our understanding of how genetic, environmental, and demographic factors influence trait diversity in diatoms, or indeed the stability of diatom traits within lineages, remains in its infancy. Trait diversification and local adaptation depend on the interplay of deterministic selective forces and random events, with models of marine microbes focusing primarily on the tension between adaptation of large populations to local environments versus their dispersal by currents (Ward et al., 2021). Although the role of chance events are rarely explicitly considered, fluctuations in population size are likely a common feature of evolving populations of marine microbes. In particular, diatom population sizes can fluctuate by orders of magnitude over the course of a bloom, and migration between ocean currents continually introduces small subpopulations into new environments (Behrenfeld et al., 2021; Ruggiero et al., 2017). Thus, repeated bottlenecks, where population size is suddenly and severely reduced, are likely a common feature for populations of marine microbes, including diatoms. In line with this, recent estimates of effective population sizes in diatoms, picoplankton, and cyanobacteria are on the order of 10^6^ - 10^7^(Chen et al., 2022; Krasovec et al., 2019; Mock et al., 2017), which are much smaller than their census population sizes. For example, diatom blooms can reach cell densities of 10^8^ cells/L (Liu et al., 2021). These discrepancies between effective and census population sizes are consistent with repeated selective events, fluctuations in population size, and frequent population subdivisions.

Experimental studies in microbial and viral populations have shown that regular population bottlenecks can profoundly affect patterns of adaptation (LeClair and Wahl, 2017). Population bottlenecks transiently reduce the supply of beneficial mutations, which can reduce total fitness gain during adaptation (Schoustra et al., 2009). However, bottlenecked populations can also better explore a higher number of alternate adaptive solutions in cases where several high-fitness phenotypes exist (Windels et al., 2021), which results in more genetic and phenotypic diversification over repeated rounds of adaptation. This broader exploration of trait values and combinations can result in higher fitness in cases where novel interactions amongst mutations occur – in these cases, more diverse, and possibly higher-fitness genotypes, may evolve from lower-fitness intermediates (Salverda et al., 2017). In contrast, in large populations with higher mutational supplies, these lower-fitness intermediates are rapidly outcompeted by the fittest possible initial mutant, which leads to fast and repeatable adaptation. Similarly, in simple environments with only a single adaptive solution, adaptation in small populations is more constrained than in large populations because deterministic adaptation rapidly locates the only available solution. However, smaller populations can better explore different adaptive solutions than can larger populations in complex environments characterized by a multiplicity of possible adaptive trajectories and endpoints. This is because smaller populations evolve less deterministically than larger ones (Jain et al., 2011; Rozen et al., 2008). Globally distributed, bloom-forming marine microbes, like diatoms, provide a compelling case to examine how trait diversity is impacted by demographic changes associated with chance environmental (e.g., advection) or ecological (e.g., blooms) events.

To investigate the impact of population bottlenecks on diatom adaptation in constant environments (i.e. absence of environmental change), we subjected six diatom strains to a series of population bottlenecks (Fig. 1) and examined shifts across the pan-transcriptome with and without bottleneck passaging. During the bottleneck phase, or “reduced selection” (RS), population growth rates decreased through time as expected due to the accumulation of deleterious mutations. At the end of the RS phase, population growth rates of viable populations were reduced by an average of 45% compared to ancestral population growth rates (Fig. S1). Following the RS phase, populations were then propagated in batch culture with large transfer sizes (i.e. back-selected populations) until population growth rates had stabilized (Fig. S2). Transcriptomes were then generated across populations. See Methods for full experimental design and Hinners et al. (2022) for in-depth analysis on whole-cell level trait changes (Hinners et al., 2022). This experimental design allowed us to study transcriptional divergence following fitness recovery, where populations had been knocked off their adaptive peaks through the accumulation of deleterious mutations during repeated bottlenecks. During fitness recovery, populations could then move towards new adaptive peaks if such peaks exist and are accessible, thereby increasing diversity within starting genotypes.

**Fig. 1.**
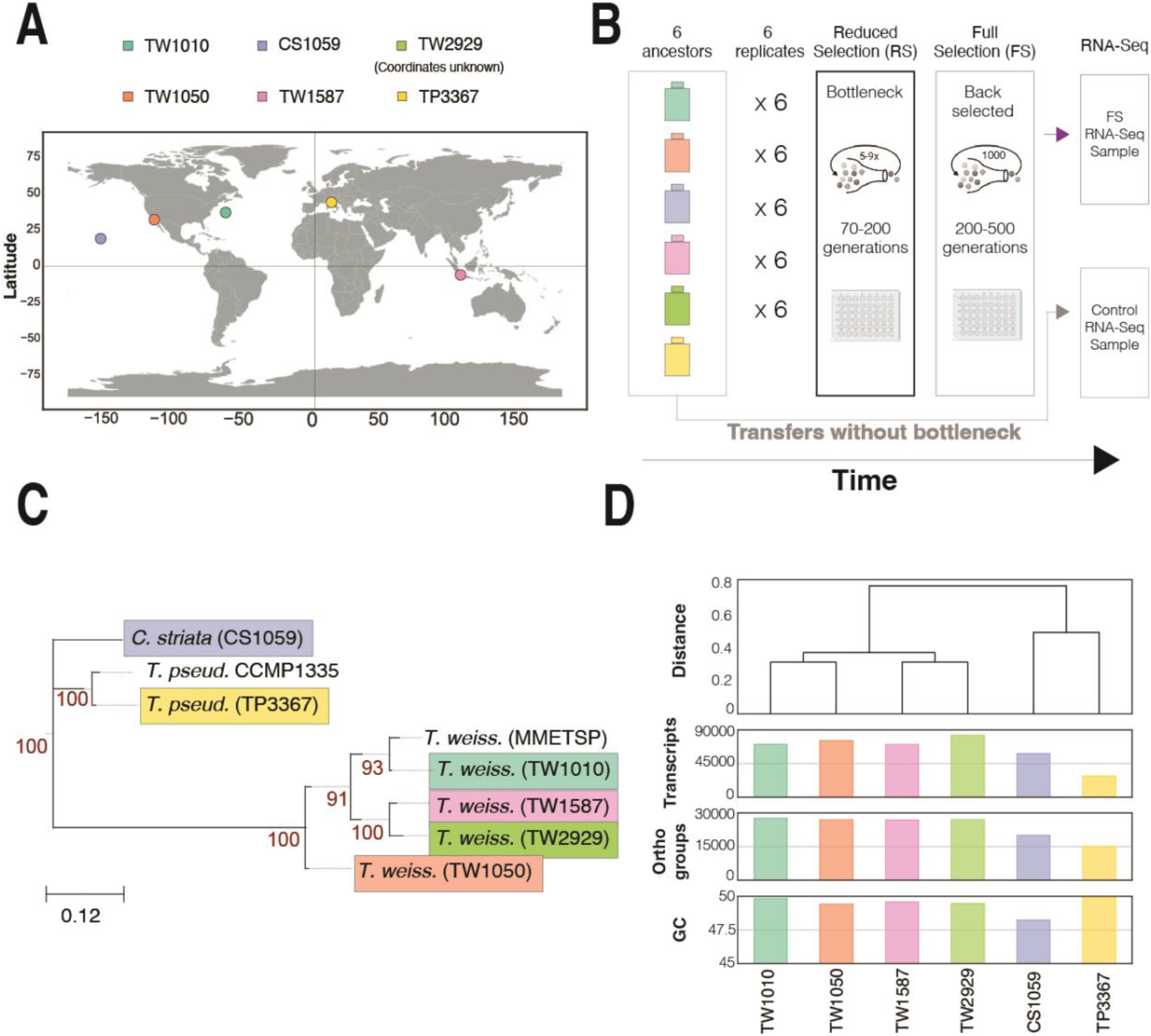
Strain information and experimental design. A) Geographic coordinates of study strains. Strain TW2929 did not have coordinates provided. (B) Experimental design where 6 ancestral strains with 6 replicates each were bottlenecked and then back-selected prior to RNA-Seq; (C) Phylogenetic tree of diatoms in study and in MMETSP (<u>Keeling 2014</u>) based on concatenated alignment of BUSCO *Protista_83.hmm* marker set. See Fig. S3 for uncollapsed tree; (D) Hierarchical clustering of jaccard distance based on shared orthogroups for study strains.

The transcriptomes of the back-selected populations (populations passed through bottlenecked followed by recovery through full selection) were analyzed to determine whether global gene expression patterns diversified following repeated bottlenecks. Transcriptional networks of most populations returned to those of the control populations following bottlenecking and full selection recovery. However, at least 1 out of 6 replicate populations from each strain traveled to other regions of the tran-scape, demonstrating that evolving a new transcriptional network as a result of bottlenecks is possible and relatively common in this system. Strikingly, most of the transcripts driving the evolution of new transcriptional networks under these conditions were orthologs shared by a subset of strains. These orthologs resided in common, central metabolic pathways across all strains and were involved in osmoregulation, cellular stress, carbon metabolism, nitrogen metabolism, and energy metabolism. These pathways may represent universal axes of variation used when diatoms evolve in response to demographic changes.

## Results and Discussion

### Strain phylogeny and orthology

To assess the relatedness of the strains used in this study, we conducted phylogenomic analysis using a set of highly conserved diatom proteins (e.g., (Keeling et al., 2014)). Our phylogeny (Fig. 1C; Fig. S3) generally agrees with the ITS2 phylogeny in Argyle et al. 2021b (Argyle et al., 2021b) demonstrating concordance between ITS2 and multi-protein sequence conservation. We next identified global orthologs across publicly available diatom genomes and our transcriptomes, which showed that unique ortholog abundance scaled linearly with unique transcript abundance (R^2^ = 0.95; Fig. S4). Hierarchical clustering based on the presence/absence of global orthologs, resulted in two main clusters with one cluster composed of the *T. weissflogii* strains and the other of *T. pseudonana* and *C. striata* (Fig. 1d). Hence, TP3367 and CS1059 share more similar numbers of global orthologs and are more phylogenetically related to each other than the TW strains, although they were isolated from vastly different locations (Fig. 1a). These phylogenetic and ortholog differences may have ancient origins, which could have been followed by subsequent ortholog and protein sequence divergence driven by environmental divergence among these diatom taxa. For example, this similarity could be driven by adaptation to warm temperature by TP3367 and CS1059 following the evolutionary divergence of TP3367 from TW strains. However, further research is needed to understand potential reasons for differences across strains. In summary, these data from our globally distributed diatom isolates demonstrate them to be relatively diverse both in phylogeny and genome characteristics, thereby representing different relatively high-fitness phenotypic solutions (i.e. sufficiently abundant *in situ* at time of sampling) within the broader diatom fitness landscape.

### Peaks and valleys in transcriptional landscapes

Selection acts on an entire organisms. Thus to assess the impact of adaptation, one must simultaneously consider the integrated phenotype composed of numerous interdependent trait relationships (Malcom et al., 2014). Here we use transcriptome data to assess how global transcriptional relationships underlying integrated phenotypes shift in response to adaptation. Here, we define a transcriptional landscape (from here tran-scape) which is similar to a trait-scape used in previous work (Argyle et al., 2021a, 2021b; Walworth et al., 2021). The transcape assumes that the control population is well adapted with defined high-fitness peaks. When adapting to a new environment or following a population bottleneck event, genotypes or strains that have moved into fitness valleys (low growth rate phenotypes) can move between peaks to arrive at high-fitness phenotypes via adaptation, so long as they avoid extinction. The main difference between a fitness landscape and a PCA-based trait-scape or PCoA-based tran-scape is that while each location on a fitness landscape defines a unique multitrait phenotype, single locations in trait- and tran-scapes can define more than one multitrait phenotype (Walworth et al., 2021). This is because each location along each non-fitness (non-growth) axis is determined by a combination of several trait values and correlations. Here we use the transcape to analyze the location of different populations in the tran-scape following fitness recovery from relaxed selection. Fig. 2a and 2b show three different high-fitness regions (peaks) based on core ortholog expression from diverse diatom strains.

**Fig. 2.**
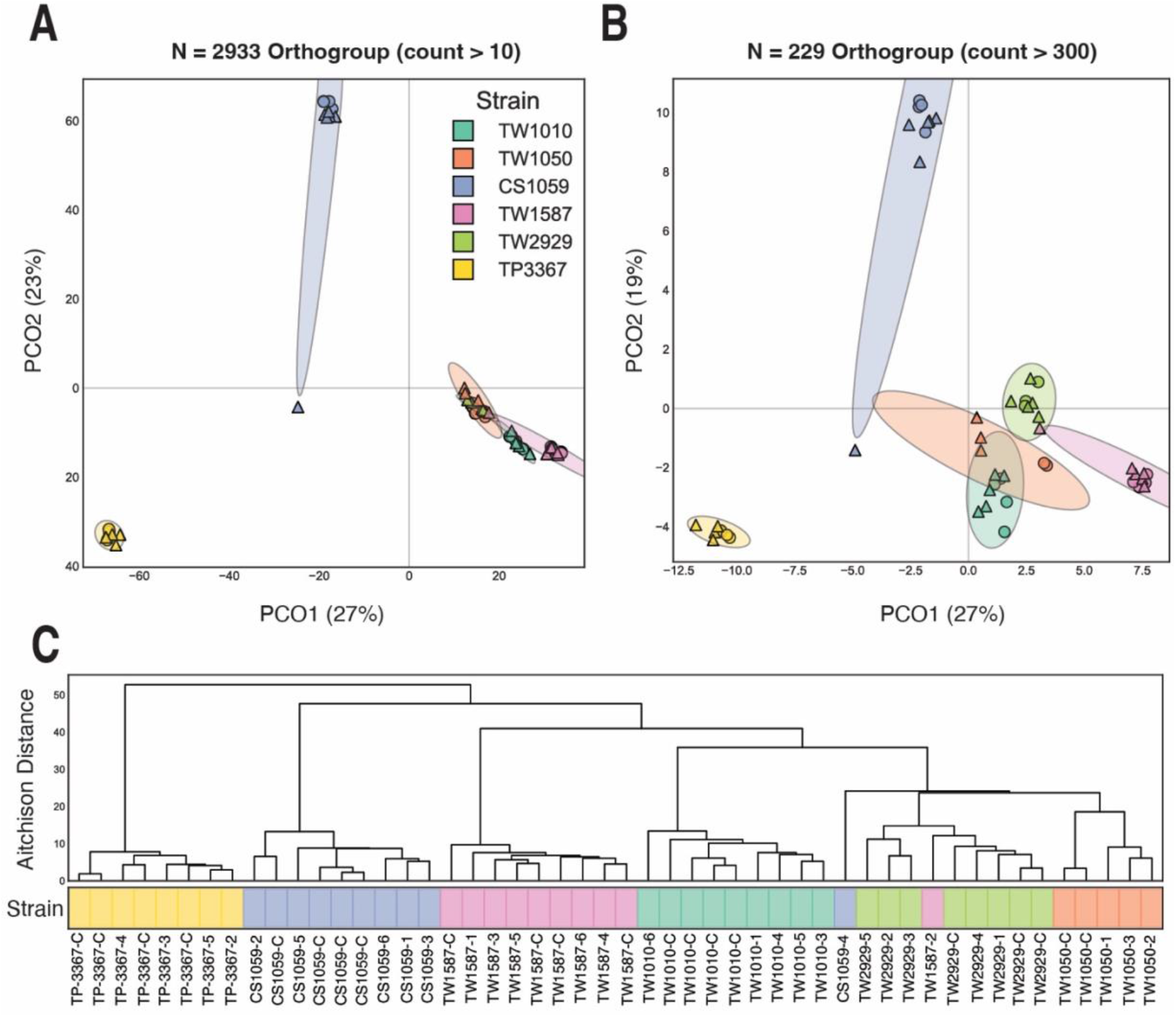
Global analysis of ortholog expression among diatom strains. Aitchison PCoA of conserved orthgroups that had a prevalence of A) 1 count per sample (N=2933 orthgroups) and B) 300 counts per sample (N=229 orthogroups); C) Hierarchical clustering using Aitchison distance of conserved orthogroup expression with at least 300 counts per sample. Circles represent ancestral controls and triangles represent bottlenecked strains followed by FS recovery (i.e. back-selected).

In general, the clustering of strains based on core ortholog expression was consistent with both the phylogenetic and global ortholog abundance clusters (Fig. 1c,d). For example, strains TW1050, TW2929, TW1587, and TW1010 core ortholog expression converge into a single high-fitness region (Fig. 2a,b). One back-selected population, CS1059-4 (Fig. 1 a,b - blue triangle in lower left quadrant; Fig. 1c), discovered an adaptive route following the bottleneck that resulted in the discovery of a peak at or near the one occupied by the TW strains. Interestingly, Hinners et al. 2022 found that most phenotypic outlier populations originated from CS1059, suggesting relatively more flexibility in trait correlations than other strains. The transcriptional divergence of CS1059-4 (Fig. 2) may indicate early molecular divergence from CS1059 replicate populations prior to more pronounced trait divergence. Next, we analyzed each strain-specific cluster to explore transcripts driving intraspecific variation at each peak.

### Strain-specific transcriptional landscapes

The tran-scape provides a means to assess whether populations returned to the ancestral high-fitness location in the tran-scape (Fig. 3, circles) or if they moved to another high-fitness location. In each strain, at least one back-selected population moved to a new high-fitness location in the tran-scape. Fig. 3 shows strain-specific tran-scapes constructed from the global transcription of each viable replicate population within each strain. The strain specific PCoA plots capture between 49-99% of total transcriptional variance. Control populations (circles) formed clusters reflecting minimal movement away from the single local high-fitness ancestral peak. TP3367 control populations exhibited more drift than other strains suggesting the ancestral peak was less stable. While this TP3367 trend is of interest, it is out of scope for this study and will be the subject of future research.

**Fig. 3.**
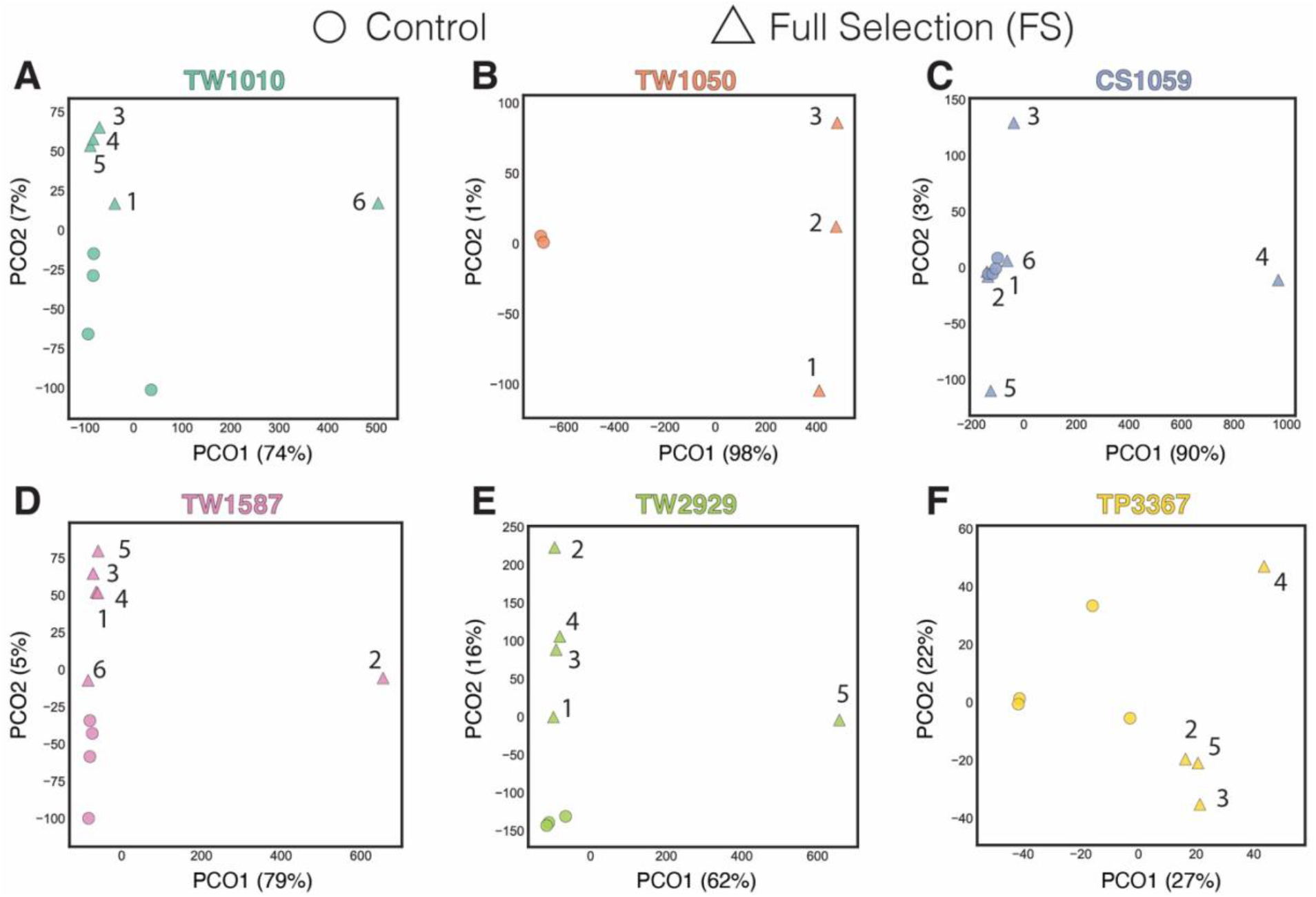
Aitchison PCoA of transcript expression for study strains. Circles represent ancestral controls and triangles represent bottlenecked strains followed by FS recovery (i.e. back-selected). Numbers next to triangles denote back-selected replicate number for each strain.

Across all strains, at least one back-selected population (triangles) found another high-fitness transcriptional peak in a region of the tran-scape not occupied by the control populations, indicated by movement along PCO1 (Fig. 3). The fact that at least one back-selected population per lineage (>17% of populations) discovered another distinct peak demonstrates a robust capability for diatoms to significantly change widespread transcriptional circuitry after a period of reduced selection. Most of the new peaks were differentiated along PCO1 – the axis which captured the majority of the variance in the ancestral populations. This suggests that, even when a new peak is found, that the new variation is captured within the ancestral tran-scape. While most of the movement occurred along PCO1, transcripts driving differentiation along PCO2 could have biological significance in generating phenotypic differences, but this is not explored further in this study.

Some strains were more stable than others. Specifically, most back-selected populations from strains CS1059, TW1587, and TW1010 returned to the ancestral peak region. Conversely, all back-selected populations from strain TW1050 migrated to an alternative high-fitness peak, with none returning to the ancestral peak region. This suggests that this lineage could be particularly prone to transcriptional diversification in the face of population size fluctuations, or that the standard culture collection isolate of this strain has a phenotype that corresponds to a local optimum that is less fit than other accessible local optima. For all strains, more fitness peaks may be identified if more replicate populations were generated and sequenced. Notably, TP3367 control populations exhibited considerable intraspecific transcriptional variation without bottlenecking (Fig. 3f, circles), which could indicate that TP3367 was adapting to general lab conditions. Additionally, less variation overall was associated with the PCO axes of this strain. Hence, we cannot be sure that diversification of back-selected populations was only due to bottlenecks. So, while TP3367 back-selected populations did seem to find two new defined peaks with TP3367-4 residing in one and TP3367-2, TP3367-3, and TP3367-5 residing in the other along PCO2 (22% explained variance), we are cautious in our interpretation due to the variability in control populations.

We next investigated if these patterns were primarily driven by the expression of the representative pangenome (global ortholog transcription) or by strain-specific genetic elements. To do this, we reconstructed the tran-scapes using only the global orthologs (Fig. S5), which yielded relatively congruent positions of controls and back-selected populations compared to tran-scapes constructed from all transcripts (Fig. 3). Fig. S5 demonstrates that diatom strains can readily rearrange global transcriptional relationships across core components of the genome following bottleneck events. These data suggest strain-specific differences in the stability of local adaptation (control peaks) with the majority of back-selected populations in some strains returning to the ancestral peak (e.g., CS1059) while none for other strains (e.g., TW1050). We next investigated what transcripts and pathways strains used to move to new high-fitness peaks.

### Identifying transcripts driving the discovery of new peaks

We first conducted pathway enrichment analysis (see Methods) on transcripts loaded onto respective strain-level PC axes to identify significantly overrepresented pathways (Supp. File 1). These analyses revealed enrichment of numerous central metabolic pathways like carbon fixation, pyruvate metabolism, glycolysis/gluconeogenesis, amino acid metabolism, porphyrin and chlorophyll metabolism, and pentose phosphate metabolism. The overrepresentation of these pathways is consistent with the fact that the majority of transcripts driving new peak discovery are orthologs (see below). Next, we identified specific transcripts in these pathways.

To examine which transcripts were most associated with the exploration of the transcape through the discovery of new high-fitness locations (Fig. 3), we analyzed transcripts harboring the largest PCoA loading values (both positive and negative) on PCO1 and PCO2 axes, respectively. Each transcript loading value reflects how much a particular transcript contributes to that principal coordinate axis. To do this for PCO1, for each strain we first tested different amounts of transcripts harboring the most positive loading values (e.g., n = 500, n = 1000, n = 1500, etc.) and conducted hierarchical clustering of the Euclidean distance among their CLR-transformed expression values. We did the same for transcripts with the most negative loading values. We then selected the amount of transcripts where at least one of the hierarchical dendrograms (either the positive or negative one) maintained a congruent clustering pattern as in the strain-specific, global tran-scape containing all transcripts in Fig. 3. For example, in Fig. 4a, TW1010-6 is replicate population number 6 of the back-selected TW1010 strain. TW1010-6 discovered a new peak in the TW1010 tran-scape (Fig. 4a, top panel) and formed its own expression cluster relative to the other replicate populations (Fig. 4a, bottom panel). Upon clustering different amounts of transcripts harboring the most negative (Fig. 4b, upper left plot) and most positive (Fig. 4b, upper right plot) loading values on PCO1, we observed that at 2000 transcripts (n = 4000 total transcripts analyzed for PCO1), at least one of the dendrograms (Fig. 4b, upper panels) reflected an analogous clustering pattern as those with all transcripts (Fig. 4a, bottom panel). In this case, the 2000 most positive loading values (Fig. 4b, upper right plot) reflected the most defined, similar pattern relative to the global tran-scape pattern (Fig. 4a, bottom plot) followed by the 2000 most negative loading values (Fig. 4b, upper left plot). Clustering beyond 2000 transcripts introduced less consistent clustering patterns relative to the global transcriptional plots indicating a greater inclusion of transcripts that did not strongly contribute to the segregation of TW1010-6 in its strain-specific tran-scape (Fig. 4a, upper panel). The strong contrast in expression values observed in the resulting transcriptional dendrograms (Fig. 4b, purple = higher relative expression and blue = lower relative expression) further reflects these most extreme loading values. Here, the most negative loading values correspond to reduced transcript levels in TW1010-6 relative to other TW1010 populations while the most positive loading values correspond to greater relative transcription. We then tested the 2000 most positive and negative loading values for all other strains for PCO1, which also yielded analogous clusterings to their respective strain-specific, global transcapes (e.g., Fig. 4c,d).

**Fig. 4.**
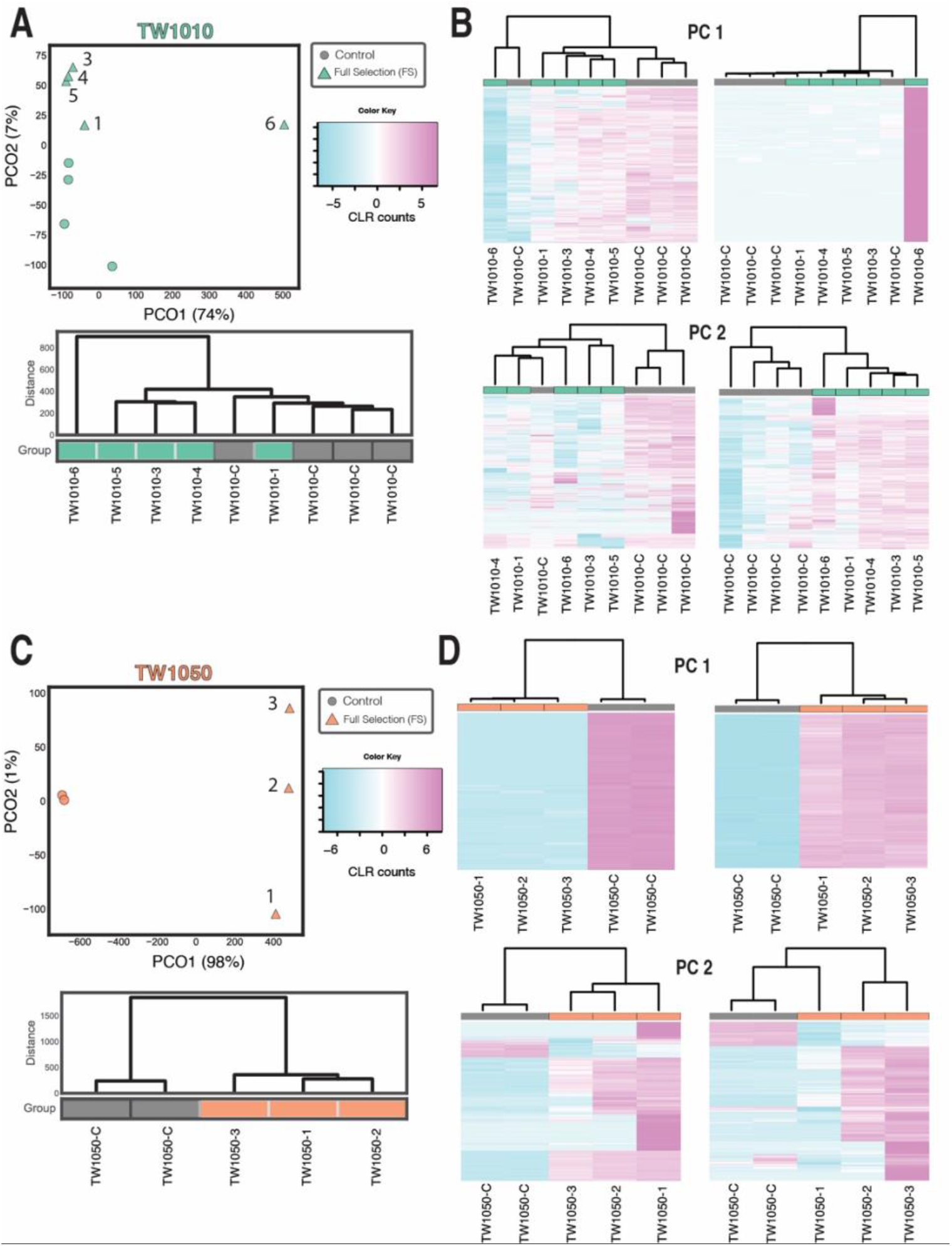
Representative analysis of transcripts driving the discovery of new peaks TW1010 and TW1050. A) Upper and lower panels are the TW1010 PCoA plot and hierarchical dendrograms of CLR-transformed transcript counts, respectively B) Upper row contains hierarchical clusters of TW1010 CLR-transformed transcript counts of the 2000 most negative (left plot) and 2000 most positive (right plot) PCoA transcript loading values for PCO1. The lower row contains the same clusters except for PCO2. C) Upper and lower panels are the TW1050 PCoA plot and hierarchical dendrograms of CLR- transformed transcript counts, respectively D) Upper row contains hierarchical clusters of TW1050 CLR-transformed transcript counts of the 2000 most negative (left plot) and 2000 most positive (right plot) PCoA transcript loading values for PCO1. The lower row contains the same clusters except for PC2 using the 1000 most positive and 1000 most negative loading values.

Due to this consistency and to also enable general comparisons across strains, we moved forward with the 2000 most positive and most negative loading values for all strains along the PCO1 axis for subsequent analyses (n = 4000 total transcripts analyzed for PCO1). These transcripts represent potential transcriptional and metabolic pathways that diatoms use to explore their transcape and to discover new peaks. We conducted the same analyses for PCO2 for all strains and found that the 1000 most positive and negative loading values (n = 2000 total) generally represented analogous clustering relative to the strain-specific, global transcape along PCO2 (e.g., Fig. 4b and 4d, bottom two plots). This reduced number of transcripts is consistent with the fact that less variance is explained on PCO2 than on PCO1, such that departures from control expression values or patterns may not represent peaks that differ very much from the ancestral one. All of these CLR- transformed transcripts harboring the most extreme loading values per PCO per strain can be found in Supp. File 2.

Fig. 4 shows two representative cases where the TW1010-6 back-selected population discovered a novel peak whereas all other back-selected populations returned to a similar region as the TW1010 controls occupy. While TW1010-1, TW1010-3, TW1010-4, and TW1010- 5 formed a cluster along PCO2 (Fig. 4a, PCoA plot; Fig. 4b, bottom right plot), it is unclear if this is a clearly defined peak distinct from the control peak due to the low variance explained on PCO2. In contrast, all back-selected replicates of strain TW1050 migrated to a single new peak along PCO1, which explains >93% of the variance (Fig. 4c,d). Hence, the TW1050 control peak may represent a less stable phenotype than the TW1010 control one. Due to the low amount of explained variance on the TW1050 PCO2 axis, no new peaks were obvious along PCO2. Finally, of these transcripts with the most extreme loading values, the vast majority represented global orthologs that were shared by some but not all strains (Fig. S6). This demonstrates that the majority of transcriptional changes associated with diversification within an environment are central components of the diatom pangenome. Other than universally expressed core orthologs across all diatoms (e.g., Fig. 2), many of these orthologous sequences are not necessarily shared by all diatom strains but rather select ones. Taken together, these data show that exploration of the tran-scape is not primarily driven by strain-specific transcripts, but through changes in widely shared pathways.

### Metabolic transcripts associated with new peak discovery

To examine the most influential metabolic transcripts driving the movement of certain replicate populations towards new peaks in this experiment, we focused on transcripts harboring the most positive loading values across PCO1 for all strains (e.g., Fig. 4b and 4d, upper right plots). These positive loading values reflect transcripts with higher expression values in the back-selected replicates that found new peaks following bottlenecking (e.g., TW1010-6, TW1050-1, TW1050-2, TW1050-3, etc.). Taken together, these transcripts represent those that uniquely increased in the replicates that found a new peak relative to the control populations.

Across all strains, numerous transcripts related to stress, reactive oxygen species (ROS), carbon metabolism, and nitrogen metabolism exhibited consistently increased relative expression levels. This is consistent with trait observations from Hinners et al. 2022 (Hinners et al., 2022) where the largest overall differences between back-selected populations relative to their ancestors and controls were in levels of ROS, particulate organic carbon (POC), particulate organic nitrogen (PON), and lipid content. For example, increased expression of numerous heat shock proteins (HSPs), aldehyde dehydrogenases (ALDHs), superoxide dismutases (SOD), aconitases, and glutathione-related transcripts was observed across strains (Supp. File 2), which is consistent with prior studies in diatoms that detected upregulation of these transcripts during physiological stress (Allen et al., 2008; Lauritano et al., 2015; Wang et al., 2020). Increased transcripts of trehalose-6-phosphate synthase were detected in TW1010, TW1050, CS1059, and TW1587. Trehalose is an intermediate, disaccharide sugar that can aid in osmotic adjustment through protein stabilization. Trehalose accumulation has been observed under iron limitation in diatoms (Allen et al., 2008) and osmotic stress in red algae (Cao et al., 2020), and can signal changes in glycolytic activity. Here, we observed these shifts not as a result of environmental change but through population bottlenecks, suggesting a more fundamental role in diatom trait diversification irrespective of specific environmental factors.

All strains also exhibited increased expression of numerous nitrogen transporters involved in nitrogen acquisition but not the reductases and hydrolases involved in nitrogen assimilation. Interestingly, increased expression of only various nitrate transporters were detected in strains TW1010, TW1050, and TW1587 while only elevated transcription of ammonium transporters were observed in CS1059 and TW2929. TP3367 highly expressed both nitrate and ammonium transporters. Furthermore, numerous glutamine fructose-6-phosphate transaminases exhibited increased expression across all strains. This enzyme is responsible for the metabolic transfer of nitrogenous groups and is involved in glutamate and amino sugars metabolism. Collectively, increased transcription of core nitrogen metabolism genes, transporters of different nitrogen species, and significant differences in PON (Hinners et al., 2022) between back-selected and control populations suggest nitrogen metabolism to be a core compensatory pathway for diatoms in the face of demographic (e.g., bottleneck) or environmental change. Its modularity (Smith et al., 2019) and redundancy (e.g., urea, aminos, nitrate, and ammonia) can aid in the sustained viability of a handful of cells exploring adaptive routes in a fitness landscape. For example, the key enzymes involved in ammonium metabolism, glutamine synthetase and glutamate synthase (GS-GOGAT), are located in both mitochondria and chloroplast in diatoms (Smith et al., 2019). This suggests potentially high amounts of near-neutral diversity in nitrogen-related traits in natural populations that experience frequent bottlenecks. Hence, shifts in nitrogen metabolism here is not indicative of adaptation to any change in the availability of nitrogen but rather a fundamental adaptive strategy associated with fitness recovery. Additionally, this modularity and redundancy may also enable nitrogen metabolism in diatoms to be easily compromised in ways that do not render cells non-viable and are thus often associated with fitness valleys. As such, fitness recovery often involves re-evolving higher functioning nitrogen metabolism, which appears to be one of the traits determining the ruggedness of the fitness landscape. Hence, these data suggest nitrogen metabolism to be especially prone to diversification in the face of bottlenecks.

In terms of carbon, energy, and core metabolic pathways, numerous carbonic anhydrase (CA) and thioredoxin transcripts also exhibited elevated expression in TW1010, TW1050, TW1587, and TP3367 indicating shifts in equilibrium between intracellular CO_2_ and HCO_3_^-^. Carbonic anhydrases can catalyze the reversible interconversion of CO_2_ and water into HCO_3_^-^ and protons and play a central role in carbon acquisition (Clement et al., 2016), though the exact role is localization-dependent (Hopkinson et al., 2016). Similar to nitrogen acquisition described above, the modularity and redundancy of carbon acquisition through numerous types of CA’s may also enable certain carbon acquisition pathways to be compromised while maintaining cellular viability. Concurrent transcriptional changes to different CA’s following bottlenecks suggests flexibility of CA-associated carbon acquisition to be an adaptive strategy during fitness recovery. Increased transcription of cytosolic malate dehydrogenase was observed in TW1010, TW1050, CS1059, TW1587, and TP2929 and is central to both the TCA cycle and gluconeogenesis. Numerous transcripts of clathrin subunits were also observed. Clathrin is a structural protein that helps deform membranes to facilitate invagination of molecules into vesicles (i.e. clathrin-mediated endocytosis). Although not highly expressed in other eukaryotes, it was found to be the 6th most abundant protein in the *T. pseudonana* proteome and plays central roles in nutrient acquisition, vesicle transport, and segregation of organelles (Nunn et al., 2009). Finally, elevated transcripts of fucoxanthin chlorophyll proteins (FCP) and other light-harvesting photosystem genes were observed (Supp. File 2). FCPs make up the key molecular complex performing light-harvesting in diatoms (Gelzinis et al., 2015) and may be fundamental to light-derived energy generation in response to significant demographic change. Changes in expression and trait values of these critical pathways have been primarily observed as a result of environmental change (e.g., (Allen et al., 2008; Bender et al., 2018; Bertrand et al., 2012; Mock et al., 2017; Smith et al., 2019)). Here, we observe collective shifts in expression across strains as a result of demographic perturbation alone, which may indicate these pathways to be prone to rapid diversification. The fact that we observe these common shifts across disparate diatom strains suggests these pathways to be key players in diatom evolution irrespective of the type (environmental or demographic) of change being experienced. Hence, these pathways may represent fundamental diatom axes that are subject to increased selective pressure as diatoms explore their fitness landscape.

### Conclusion

Here, we demonstrate the utility of an integrated approach via generating demographic change (i.e. bottlenecking) while examining underlying global transcription across geographically disparate diatom strains. These data reveal local peaks with stable transcriptional networks and further demonstrate the ability of certain populations to find new metabolic configurations, thereby surfacing other accessible genotypic and phenotypic potential available to diatoms. The concurrent changes in both traits (Hinners et al., 2022) and transcripts across carbon, nitrogen, energy, and oxidative stress pathways indicate that they can shift in a way that is associated with lower-fitness (but viable) transitions. In other words, changing trait and transcriptional relationships is one way to descend into a non-lethal fitness valley. Additionally, during fitness recovery, these traits are part of a rugged fitness landscape where multiple solutions for high-fitness trait values and trait combinations exist, lending these traits to likely be especially involved in diversification. Finally, these peaks are connected by fitness valleys that can be traversed (growth reductions on the order of 50%). Future studies investigating inter-replicate divergence as well as the addition of more distantly related ancestral diatom strains harboring different transcriptional circuitry would shed further light on the connectivity of high-fitness peaks and low-fitness valleys. Populations are likely to descend into these valleys as a result of natural selection being temporarily relaxed during population bottlenecks, as changes were observed in these traits and transcripts repeatedly. Thus, it is reasonable to hypothesize that these traits, which are involved in fitness recovery and diversification within environments, are also involved in adaptation to new environments where a shift in environment rather than a shift in demography is responsible for putting the population into a valley. These patterns have evolutionary implications for identifying metabolic pathways used during trait diversification, including evolutionary constraints that inhibit access to certain regions of the tran-scape.

It is noteworthy that these trends occurred in nutrient-replete and generally benign conditions, which revealed the fundamental flexibility of certain metabolic pathways. Future studies can explore the conservation of these trends when fitness recovery occurs under more stressful or realistic conditions, where only a subset of the observed pathways may be able to evolve. Broader investigation of these axes in other phytoplankton taxa can yield insights into photoautotrophic evolutionary potential that may help tease out general cellular responses to selection vs those that are environment or genotype specific. Understanding these general constraints adds critical knowledge to genotypic and phenotypic limits of globally different taxa as they experience local demographic and environmental selection. Importantly, the discovery of new fitness peaks following population bottlenecking provides an evolutionary mechanism that can help explain global patterns in local adaptation and microdiversity independent of environmental selection. Hence, demographic changes lead to lineage diversification that can expand the breadth of genotypic and phenotypic space available for selection to changing conditions.

## Methods

### Diatom cultures

Six strains of *Thalassiosira sp*. from the Provasoli-Guillard National Center of Marine Phytoplankton (NCMA, formerly known as the CCMP, https://ncma.bigelow.org/) culture collection were used: CCMP 1010, 1050, 1059, 1587, 2929 and 3367 (Table S1). Extensive trait and phenotypic characterization of these strains are described in (Argyle et al., 2021a, 2021b; Hinners et al., 2022).

### Evolution experiment

#### Overview

A detailed description of the experimental design (Fig. 1B) can be found in (Hinners et al., 2022). Here we only analyzed populations back-selected in the ancestral temperature 20°C following the bottleneck while Hinners et al. 2022 subjected back-selected populations to two different temperatures (20°C and 24°C). The experiment was divided into two main phases where phase I consisted of an initial 3-month long reduced-selection (i.e. bottleneck) phase (corresponding to 70-200 generations) followed by an 8-month full selection (i.e. back-selected) phase (200-500 generations). Six replicates of each of the six ancestral populations (n = 36 cultures) were subjected to this bottleneck phase at 20°C followed by the back-selected phase also at 20°C. Trait measurements including growth rates were performed at the beginning of the experiment (Fig. S1, filled gray circle) and throughout the bottleneck and back-selected phases.

#### Reduced Selection phase

During the reduced selection (RS) phase, bottlenecks were induced every 7 days by transferring ~8 cells per replicate to new medium in order to reduce the efficacy of natural selection and allow divergent lower-fitness phenotypes to evolve. As growth rates decreased through time, we extended the bottleneck period to every 14 days towards the end of this phase resulting in an average of 18 generations. All populations were bottlenecked at the same time as long as replicates had reached a minimum cell concentration of 2000 cell/mL. If cell concentrations were lower, cultures were instead diluted to 500 cell/mL to allow for population recovery before a new bottleneck. Bottlenecks were repeated 5-9 times depending on the genotype corresponding to a total RS phase length of 3 months (70-200 generations), depending on population growth rates. Towards the end of this phase, some population growth rates decreased to a degree where growth was no longer observed. In these cases, previously saved transfers were used to induce a new bottleneck. Growth rates were monitored via in vivo fluorescence, and at the end of this phase, population growth rates were reduced by an average of 45% compared to ancestral growth rates (Fig. S1)

#### Full Selection phase

During the full selection (FS) phase, back-selected populations were propagated in batch culture with transfer sizes of 1000 cells every 7 days in the ancestral environment. Populations were transferred ~25 times corresponding to 200-500 generations. Maximum growth rates were measured every 5-10 transfers to monitor fitness recovery, and before termination of this phase, growth rates were measured over 4 transfers (4 weeks) to ensure population growth rates had stabilized, indicating populations were on or near a high-fitness peak (Fig. S2).

#### Growth rate

Growth rates were measured through daily *in-vivo* fluorescence with a Tecan Spark plate reader (excitation: 455 nm, emission: 620 nm)(Hinners et al., 2022). Exponential growth rates were calculated for each time step as follows:

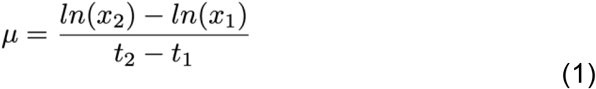

Maximum growth rates were determined over 4 consecutive time steps. During the bottleneck phase, growth rates were determined on single replicates per population. Final growth rates were determined from 3 replicates per population.

### Phylogenomics and geographic visualization

Phylogenomic trees were inferred with *IQTREE2* (Minh et al., 2020) using concatenated protein alignments constructed via *hmmsearch* for marker detection, *MUSCLE* (Edgar, 2004) for marker protein alignment, and *ClipKIT (Steenwyk et al., 2020*) for alignment trimming. The concatenated alignments were based on the BUSCO *Protista_83.hmm* (Simão et al., 2015) marker set available through *Anvi’o (Eren et al., 2015)* using the suggested E-value noise cutoff of 1e-25. Phylogenomic trees were analyzed and visualized using ETE 3 (Huerta-Cepas et al., 2016) in Python.

Geographic coordinates in relation to strain origin were processed using *GeoPandas* (https://github.com/geopandas/geopandas), *GeoPy* (https://github.com/geopy/geopy), and *Matplotlib* (https://github.com/matplotlib/matplotlib).

### Transcriptome assembly, gene modeling, and orthology

Sequence reads were quality-controlled using *KneadData* (Beghini et al., 2021) with the GRCh38.p13 human genome as a reference for potential decontamination. This methodology yielded transcriptomes with depths between 3,433,160 and 26,023,146 reads mapping between 14,761 and 55,849 unique transcripts. Refer to Supp. File 1 for richness and depth statistics per strain. De novo transcriptomes were grouped by strain (e.g., TW1010 ancestors and bottlenecks) and co-assembled using *rnaSPAdes (Bushmanova et al., 2019*).

Following the protocol detailed in Santoro et al. 2021 (Santoro et al., 2021), we used *TransDecoder* (https://github.com/TransDecoder/TransDecoder) for gene modeling in a multistep process to minimize false positives. In particular, we used the following procedure: (i) *TransDecoder.LongOrfs*, with transcript-to-gene mappings assigned by *rnaSPAdes*, to generate putative open reading frames (ORFs); (ii) *hmmsearch* (Eddy, 2011) to identify protein domains using the *PFAM* v33.1 and *TIGRFAM* v15.0 databases; (iii) *Diamond blastp* (Buchfink et al., 2021) against all *Bacillariophyceae* (diatoms) genomes available in NCBI (GCA_000149405.2, GCA_000150955.2., GCA_000296195.2, GCA_001750085.1, GCA_002217885.1, and GCA_900660405.1); and (iv) *TransDecoder.Predict* with the putative ORFs from (i), the protein domains from (ii), and the alignments from (iii) using the --single_best_only argument. Genes were annotated by best-hit *Diamond* blastp alignment to NCBI’s non-redundant protein database (accessed on v2021.08.03).

Orthogroups were identified using OrthoFinder (Emms and Kelly, 2019) with the high-quality proteins generated from our *TransDecoder* procedure and all of the *Bacillariophyceae* proteins listed previously. Consensus annotations for orthogroups were assigned by using the most common organism-agnostic annotation within the grouping using *UniFunc*, a natural language processing software developed for bioinformatics (Queirós et al., 2021).

### Pathway Enrichment Analysis

We performed KEGG pathway enrichment analysis on each strain using the *GSEA’s Prerank* rank module (Subramanian et al., 2005) via the *GSEApy* Python package (Fang et al., 2021). To prepare the data for pathway enrichment, we aggregated the counts for transcripts by their BRENDA enzyme representative (e.g., EC:1.1.1.1) and identified conserved enzymes that had at least 300 counts in each sample which were later used for pathway enrichment. The enzyme counts matrix (i.e., sample vs. enzymes) was CLR transformed followed by Euclidean distance (i.e., Aitchison distance) and PCoA was performed. The PCoA loadings of the conserved enzymes were used as feature ranks (e.g., weights) in the *Prerank* module using min_size=5 and permutation_num=1000 parameters. Enriched pathways were considered significant if FDR < 0.25 which is recommended by the *GSEA* documentation.

### Transcript analysis

Taking a compositional approach, we used the centered log-ratio (CLR) transformation on raw transcript counts by taking the log of each count and dividing by the geometric mean using the *compositional* Python package (Espinoza et al., 2020). Hierarchical clustering and PCoA ordinations was performed using *SciPy* (Virtanen *et al*., 2020) and *Soothsayer* (Espinoza *et al*., 2021) Python packages. Heatmaps were generated using the *heatmap* function in the R *stats* package (https://www.R-project.org/).

## Acknowledgements

For the purpose of open access, the author has applied a Creative Commons Attribution (CC BY) license to any Author Accepted Manuscript version arising from this submission. This research was supported by the Gordon and Betty Moore Foundation Marine Microbes Initiative (MMI 7397 SC, NL and MD) and NSF OCE-1558453 and NSF OCE- 2049299 to CLD.

## Supplementary Figures

**Fig. S1.**
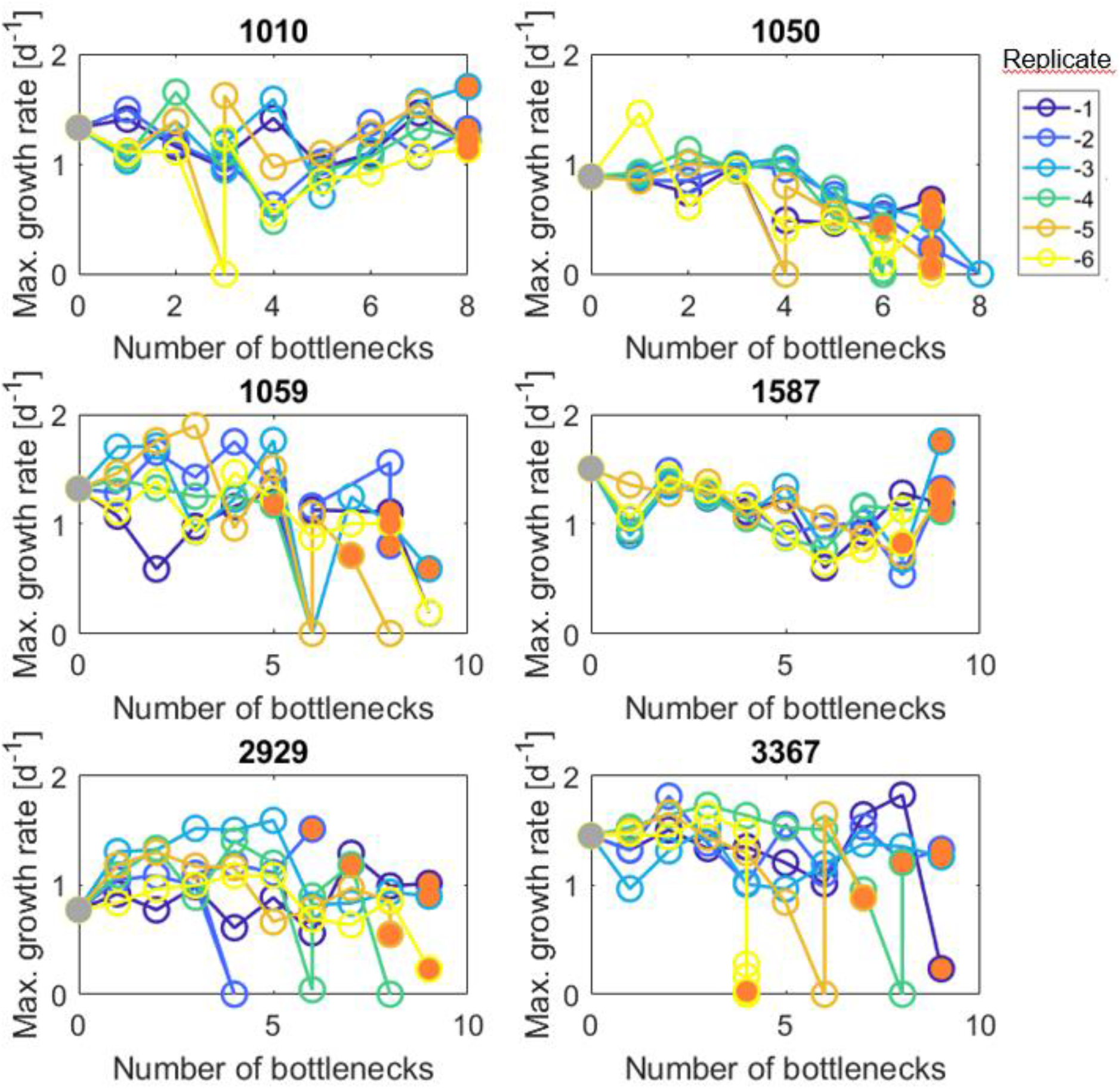
Maximum growth rates during the bottleneck (reduced selection) phase. Growth rates represent single replicates. Filled gray circles denote ancestral growth rates and orange filled circles denote final populations used for subsequent back-selected experimental phase. Vertical lines denote growth rates that decreased to such a degree that populations could not be maintained, and the bottleneck had to be repeated with a culture from the previous transfer. Strains TW1010 and TW1587 only exhibited short-term fluctuations whereas TW1050 and CS1059 consistently decreased in growth. Select single replicates of TW2929 and TP3367 responded strongly to bottlenecks through reduced growth. At the end of the bottleneck phase, average maximum growth rates were reduced by 45% relative to ancestral growth rates. Figure adapted from Hinners et al. 2022 (Hinners et al., 2022).

**Fig. S2.**
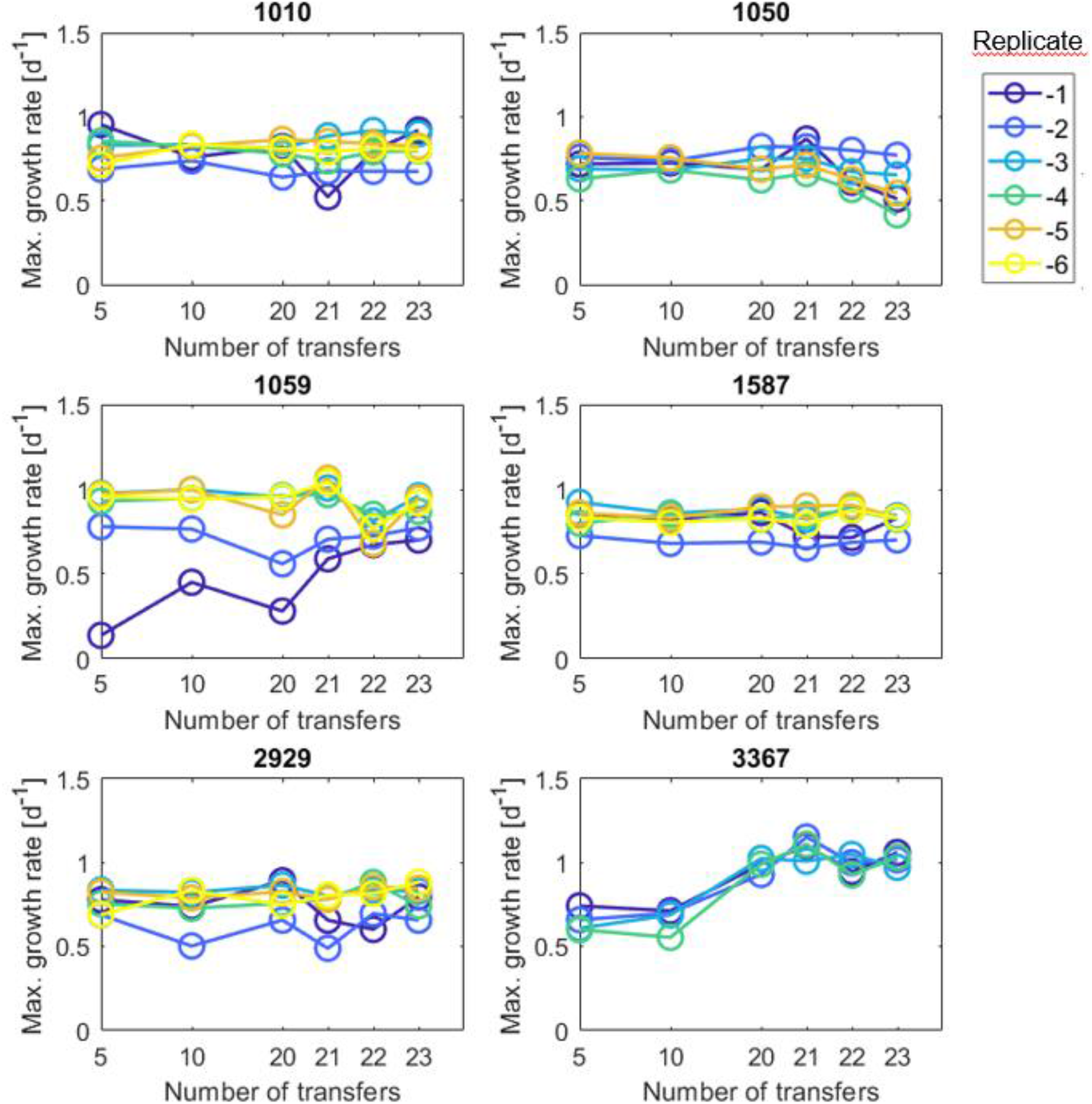
Maximum growth rates during the full selection phase. Growth rates represent single replicates. Most strains exhibited stable growth rates following the back-selected phase. During the bottleneck phase, CS1059 and TP3367 experienced strong reductions in growth rates but were able to recover over the back-selected phase.

**Fig. S3.**
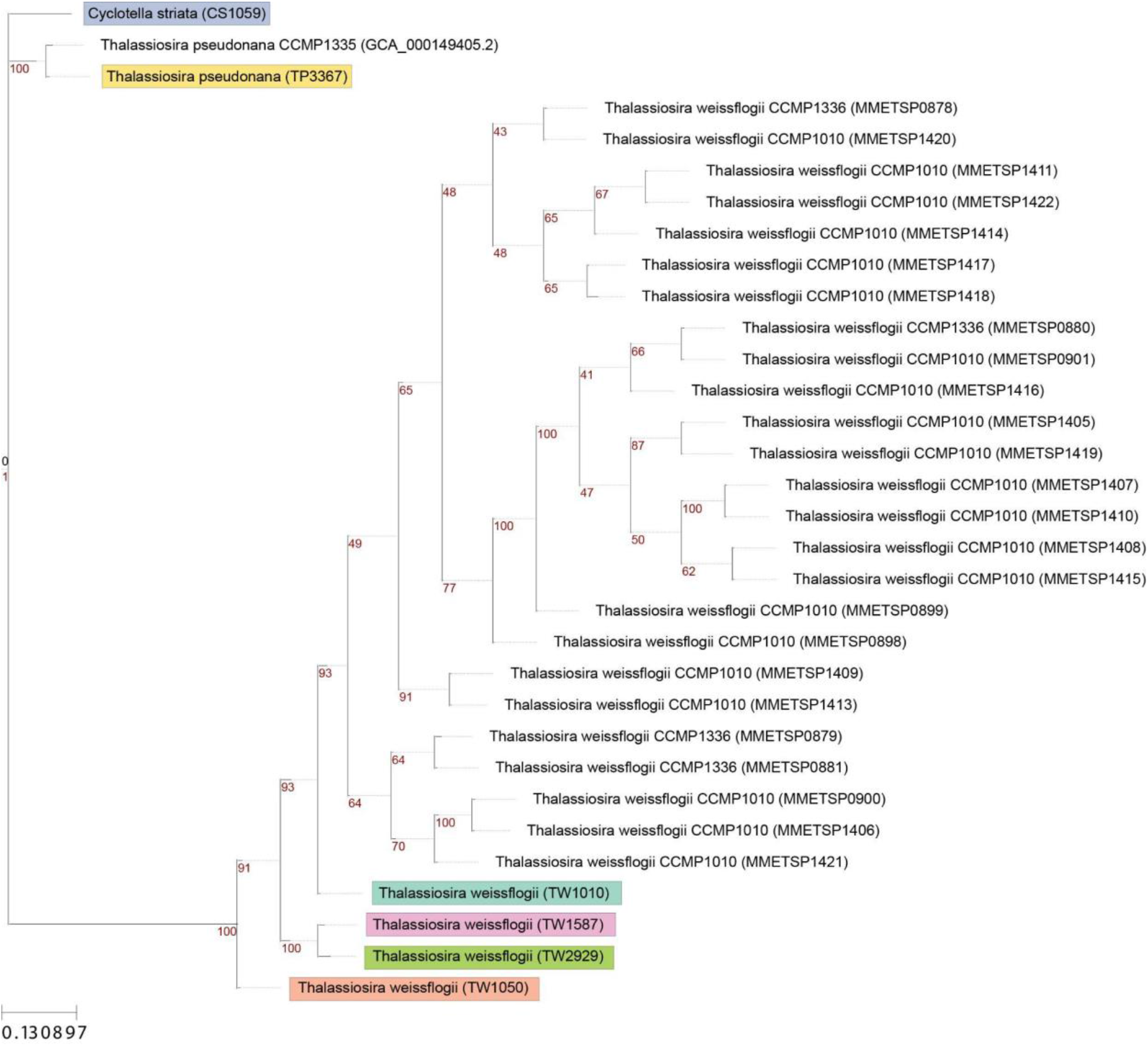
Phylogenetic tree of diatoms in study and in MMETSP based on concatenated alignment of BUSCO *Protista_83.hmm* marker set. Red numbers are bootstrap values.

**Fig. S4.**
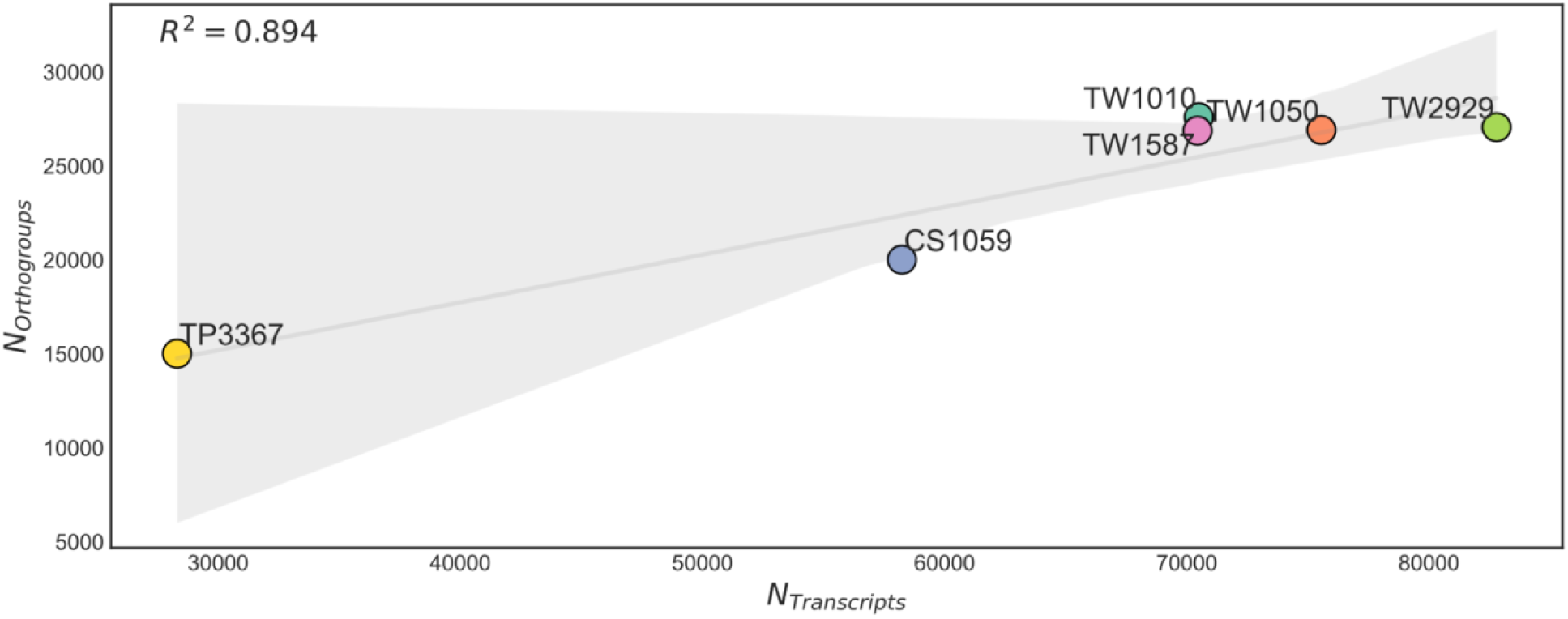
Regression plot showing linear relationship between the number of transcripts and the number of orthogroups in study strains.

**Fig. S5.**
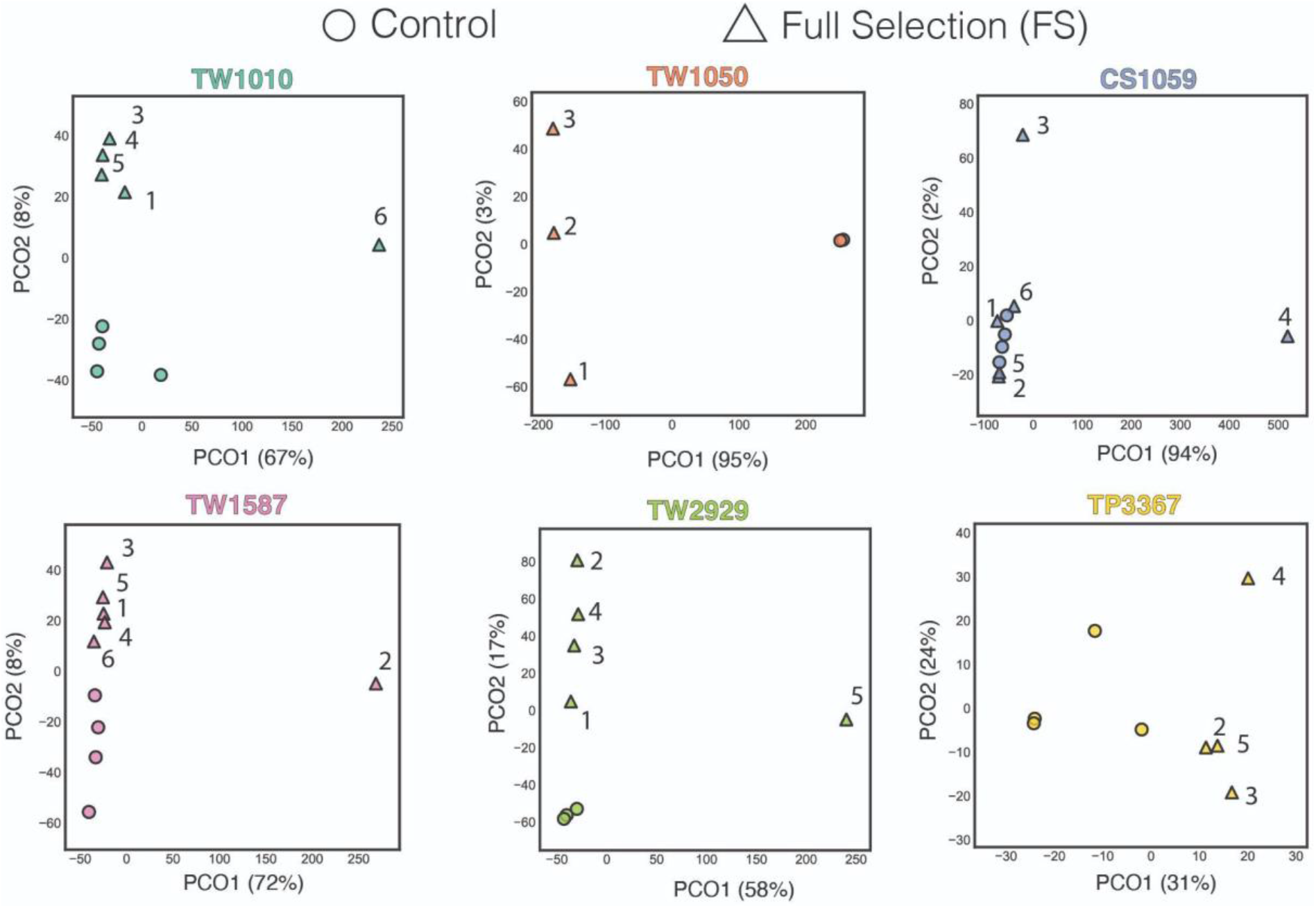
Aitchison PCoA of orthogroup expression for study strains. Circles represent control populations and triangles represent back-selected populations.

**Fig. S6.**
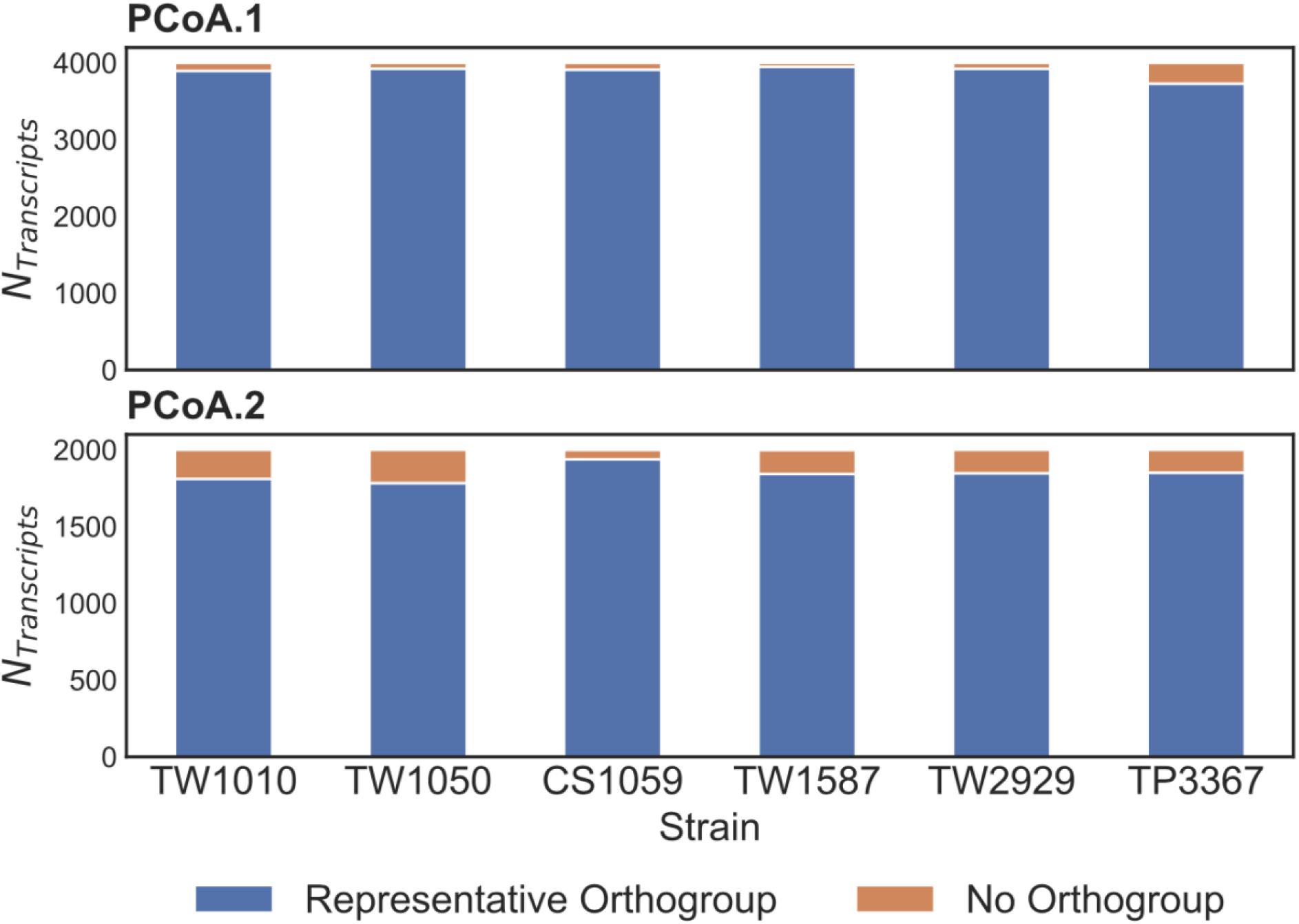
Stacked barchart of transcripts with highest loadings in PCoA including top 2000 positive loading and top 2000 negative loading transcripts across all strains. Transcript products are either represented in an orthogroup (blue) or are singleton and are not represented in an orthogroup (orange).

**Table S1.** Characteristics of strains in this study. ITS2 identification was performed by (Argyle et al., 2021). Table was adapted from (Hinners et al., 2022).

| Strain code | Strain code for this study | Collection Location | Species | ID based on ITS2 region |
| --- | --- | --- | --- | --- |
| CCMP1010 | TW1010 | 37°N 65°W, Gulf Stream, between Bermuda and New York (approx.) | *T. weissflogii* | *T. weissflogii* |
| CCMP1050 | TW1050 | 32.966°N 117.251°W, Del Mar Slough, California USA | *T. weissflogii* | *T. weissflogii* |
| CCMP1587 | TW1587 | 6.08° S 106.79°E , Jakarta Harbor, Indonesia (approx.) | *T. weissflogii* | *T. weissflogii* |
| CCMP1059 | CS1059 | 19.665°N 156.034°W Aquaculture Pond, Oahu, Hawaii USA (approx.) | Thalassiosira sp. | *Cyclotella striata* |
| CCMP2929 | TW2929 | Unknown | Thalassiosira sp. | *T. weissflogii* |
| CCMP3367 | TP3367 | 44.9335°N 12.7005°E North Adriatic Sea | *T. pseudonana* | *T. pseudonana* |

